# Uptake of Prochlorococcus-derived metabolites by *Alteromonas macleodii* MIT1002 shows high levels of substrate specificity

**DOI:** 10.1101/2025.01.10.632383

**Authors:** Kathryn H. Halloran, Rogier Braakman, Allison Coe, Gretchen Swarr, Melissa C. Kido Soule, Sallie W. Chisholm, Elizabeth B. Kujawinski

## Abstract

Seawater contains an abundance of small biomolecules, or metabolites, that are highly labile components of dissolved organic matter (DOM). Marine microbes interact by exchanging metabolites, thus shaping marine microbial ecology, DOM composition, and global carbon cycling. To better constrain one set of microbe-metabolite interactions, we cultured the marine gammaproteobacterium *Alteromonas macleodii* MIT1002 on a range of compounds excreted by a sympatric cyanobacterium, *Prochlorococcus*. *Alteromonas* could assimilate the branched chain amino acids leucine, isoleucine, and valine, as well as 3-methyl-2-oxobutanoic acid (a branched chain ketoacid intermediate of valine metabolism), but not thymidine, kynurenine, 4-hydroxybenzoic acid, or the other branched chain ketoacids. The assimilation of branched chain amino acids indicates that *Alteromonas* can metabolically process each corresponding ketoacid, suggesting that transporter specificity underlies the observed substrate specificity for 3-methyl-2-oxobutanoic acid. These experiments show that even subtle changes in chemical structure can result in different microbial interactions and different fates for dissolved metabolites.

**Significance Statement:** Microbial interactions with dissolved organic matter are important controls on the marine carbon cycle. Dissolved organic matter is often considered in bulk, which leaves the specificity and nature of these interactions poorly constrained. Here we show that microbe-molecule interactions can be highly specific, distinguishing between molecules that are structurally and biochemically similar. This implies that small changes in this pool of carbon could have large impacts for the overall system function, and that measuring this pool of carbon with molecular-level resolution is important to characterizing microbe-molecule interactions. We further explore the mechanism underlying the observed substrate specificity and suggest that it is caused by transporter selectivity, meaning the ability of these microbes to selectively uptake specific dissolved organic molecules.

## Introduction

Dissolved organic carbon (DOC) makes up 662 Pg of carbon in the ocean, a reservoir of carbon which is comparable to atmospheric carbon dioxide (Hansell et al., 2009). Marine microbes influence the composition and flux of DOC, and molecules in this carbon pool mediate microbial interactions. Elucidating these microbe-molecule connections is therefore crucial to understanding both microbial ecology and global carbon cycling (Moran, Kujawinski, et al., 2022). While most studies have examined the response of microbes to bulk dissolved organic matter (DOM) or broad molecular subsets of DOM (Givati et al., 2024; Liu et al., 2020; Moran, Ferrer-González, et al., 2022; Pedler et al., 2014), the specificity of interactions between microbes and molecules within DOM remains an open question. For example, recent work has shown that diverse heterotrophic bacteria preferentially use different classes of organic carbon in a way that cannot be simply predicted by phylogeny (Forchielli et al., 2022). Similarly, bacterial growth efficiency (BGE), or the fraction of carbon incorporated into biomass relative to carbon uptake, is expected to vary with both microbial taxa and organic substrate (Saifuddin et al., 2019). This heterogeneity implies that a small change in either the microbial community or the composition of DOM could significantly impact the ecology and carbon flux of marine environments.

The picocyanobacterium *Prochlorococcus* is the dominant phototroph in subtropical oligotrophic oceans (Partensky et al., 1999), responsible for fixing an estimated ∼4-8 Pg of carbon per year (Flombaum et al., 2013; Letscher et al., 2023). An estimated 2-25% of this carbon is exuded as DOC by actively growing cells (Bertilsson et al., 2005; López-Sandoval et al., 2013). The quantity and composition of this exudate varies with strain and environmental conditions, providing a complex set of substrates which can mediate interactions between *Prochlorococcus* and other marine microbes (Becker et al., 2019; Biller et al., 2016; Calfee et al., 2022; Kujawinski et al., 2023; Ofaim et al., 2021; Sher et al., 2011; Szul et al., 2019).

*Alteromonas macleodii* (hereafter referred to as *Alteromonas*) is a model marine copiotrophic microorganism (Pedler Sherwood et al., 2015) often isolated from *Prochlorococcus* enrichments (Aharonovich & Sher, 2016; Biller et al., 2015, 2016). This heterotrophic microorganism has been used to probe microbial sinks of DOC, as *Alteromonas* cultures can decrease the concentration of bulk DOC as effectively as a bulk microbial consortium (Pedler et al., 2014). In co-cultures, *Alteromonas* lives mutualistically with *Prochlorococcus*, easing oxidative stress for the phototroph while benefiting from *Prochlorococcus*-derived DOC (Biller et al., 2016; Morris et al., 2008, 2011, 2012). Despite the copiotrophic lifestyle of *Alteromonas*, individual strains use different subsets of organic molecules, highlighting substrate specificities that likely reflect the environments from which each strain was isolated (Ivars-Martinez et al., 2008; Koch et al., 2020; Manck et al., 2020).

Recent work quantified *Prochlorococcus*-derived metabolites (Kujawinski et al., 2023), which has provided new insight into the microbial sources of DOM and the specific molecules which make up this pool of carbon. The fate of those molecules and their effect on microbial ecology remains unconstrained. Here we investigate whether *Alteromonas* can grow on various *Prochlorococcus*-derived metabolites and undercover evidence for high degrees of substrate specificity.

## Methods

### Strain and Media

All experiments were conducted with *Alteromonas macleodii* MIT1002, a strain initially isolated from *Prochlorococcus* NATL2A cultures (Biller et al., 2015). Media were prepared by filtering (0.2 µm, Pall PES membrane filters) and autoclaving Vineyard Sound seawater, followed by amendment with ProMM nutrients (Berube et al., 2015) (Supplemental Information). Under sterile conditions, cultures were revived from cryopreserved glycerol stocks and grown in ProMM media (Supplemental Information). Cultures were transferred three times during mid-exponential growth in ProMM prior to each experiment. Reagent purity details can be found in Supplemental Information.

### Growth experiments

Preliminary experiments confirmed that *Alteromonas* could grow without exogenous vitamins, on sodium pyruvate (hereafter referred to as pyruvate) alone, and on lower carbon concentrations than present in ProMM (Supplemental Information, Figures S1, S2). To test the assimilation of metabolites known to be exuded by *Prochlorococcus*, we cultured *Alteromonas* on a series of these compounds. In each growth experiment, *Alteromonas* received either a *Prochlorococcus*-associated metabolite, pyruvate, or a mix of pyruvate and the metabolite.

Pyruvate acted as a positive control, as preliminary experiments confirmed the ability of *Alteromonas* to grow on this substrate alone. Mixes of pyruvate and the selected *Prochlorococcus* metabolite were tested, at multiple pyruvate concentrations, in case metabolite utilization was enabled when pyruvate supplemented the energetic demands of *Alteromonas*.

Metabolites tested were chosen based on their presence in *Prochlorococcus* spent media (Kujawinski et al., 2023) and included: thymidine; 4-hydroxybenzoic acid; kynurenine; the branched chain amino acids valine, isoleucine, and leucine; and the branched chain ketoacids 3-methyl-2-oxobutanoic acid (intermediate of valine metabolism), 3-methyl-2-oxopentanoic acid (intermediate of isoleucine metabolism), and 4-methyl-2-oxopentanoic acid (intermediate of leucine metabolism). Cultures were grown with 0 or 900 µM C from the *Prochlorococcus*-derived metabolite, and 0, 900, or 1800 µM C from pyruvate (totaling 0, 900, 1800, or 2700 µM C; for reference, ProMM contains 6900 µM C). If a metabolite failed to support *Alteromonas* growth, results were verified with repeat growth experiments (Figures S3, S4).

Growth experiments were conducted on black walled, clear bottom, 96-well plates using a BioTek Synergy 2 plate reader, measuring OD_600_ every 30 minutes for 48 hours. Milli-Q blanks were used in wells along the border of the 96-well plate to mitigate growth discrepancies due to evaporation in exterior wells. Additional growth discrepancies due to well position were occasionally observed on the next row or column in from the edge. If these discrepancies were observed in any single sample of an edge row or column, the entire row or column was removed from analysis for that experiment. Experiments were conducted with six replicates for each condition. After removing well plate columns or rows where replicates crashed, we were left with at least three replicates for each condition (except for the leucine growth experiment, where due to random chance the 900 µM C from pyruvate condition was left with two replicates). The average signal in the 0 µM C negative controls was subtracted from each well as a baseline correction, to account for trace seawater carbon. The carrying capacity in each well was calculated as the difference between the maximum and minimum OD_600_ observed in that well. The significance of the effect of carbon substrates on carrying capacity was calculated using one-way ANOVA with a post-hoc Tukey’s HSD test.

### Uptake experiments

To test whether *Alteromonas* could take up compounds without a concomitant increase in biomass, we measured the uptake of 3-methyl-2-oxobutanoic acid and 3-methyl-2-oxopentanoic acid during exponential growth by monitoring the change in media (i.e., extracellular) metabolite concentrations. Before uptake experiments, cultures were washed 3 times with unamended sterile seawater by centrifugation for 10 minutes at 7000 x *g*. Cultures were then grown in 100 mL of minimal media containing seawater, inorganic nutrients, and 180 µM of a selected metabolite (Supplemental Information) in 125 mL acid-washed polycarbonate bottles. Control cultures were also grown with no metabolite added. To ensure we would sample during exponential growth, one culture was grown in parallel with experimental cultures, using 3-methyl-2-oxobutanoic acid as the metabolite, and sampled repeatedly. At each timepoint, cultures and controls were sampled destructively in triplicate. 750 µL of each culture or control was fixed with paraformaldehyde and stored at -20 °C until analysis by flow cytometry (see Supplemental Methods for flow cytometry details). The remaining culture was filtered through an Omnipore (Millipore) 0.2 µm PTFE membrane filter. Filtrate was acidified to pH ∼3 using concentrated hydrochloric acid and stored at 4 °C for less than 48 hours until PPL extraction.

Acidified filtrate was extracted with 100 mg, 1 mL Bond Elut PPL cartridges (Agilent, Santa Clara, CA, USA) as previously described (Dittmar et al., 2008; Kido Soule et al., 2015). Extracts were dried on a vacufuge (Eppendorf) and reconstituted in Milli-Q water with deuterated biotin (final concentration 0.05 mg mL^-1^) and ^13^C_5_-3-methyl-2-oxobutanoic acid (final concentration 0.02 µg ml^-1^). Extracts were stored at -20 °C until analysis by liquid chromatography-tandem mass spectrometry (LC-MS/MS; see Supplemental Methods for details on LC-MS/MS).

Standard curves were constructed using at least a five-point matrix-matched standard curve which spanned the range of observed sample signal intensities. For 3-methyl-2-oxobutanoic acid, standard curves were constructed using the signal of unlabeled 3-methyl-2-oxobutanoic acid normalized to the signal of ^13^C_5_-3-methyl-2-oxobutanoic acid. 3-methyl-2-oxopentanoic acid standard curves were constructed using the signal of unlabeled 3-methyl-2-oxopentanoic without additional normalization. Measured concentrations were corrected for extraction efficiency (Johnson, Kido Soule, and Kujawinski 2017; Kujawinski et al. 2023). For 3-methyl-2-oxobutanoic acid, metabolite concentrations were normalized to cell concentrations.

The resulting per-cell bulk uptake rate was treated as a pseudo-first order process and modeled as an exponential decay function where C*_t_* is the concentration at time *t*, C_0_ is the initial concentration, and *k* is the per-cell uptake rate (Equation 1).

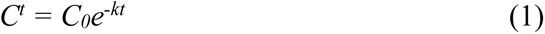

## Results

### Growth experiments

Of the metabolites tested, only the branched chain amino acids (valine, isoleucine, and leucine) and the valine precursor 3-methyl-2-oxobutanoic acid supported *Alteromonas* growth (Table 1; Figures 1 – 2, S3 – S5). In general, *Alteromonas* reached a significantly higher carrying capacity when grown with one of these four metabolites alongside pyruvate than when grown with pyruvate alone, although this trend does not hold for all four of these metabolites across all pyruvate concentrations tested (Figure 2). However, at constant carbon concentrations, carrying capacity is significantly higher on pyruvate alone than on any of these four metabolites, or on a mix of pyruvate and any of these metabolites (Figures 2A, S5 – S7). In contrast, 3-methyl-2-oxopentanoic acid, 4-methyl-2-oxopentanoic acid, thymidine, kynurenine, and 4-hydroxybenzoic acid did not support growth (Figures 1 – 2, S3 – S4)..

**Figure 1.**
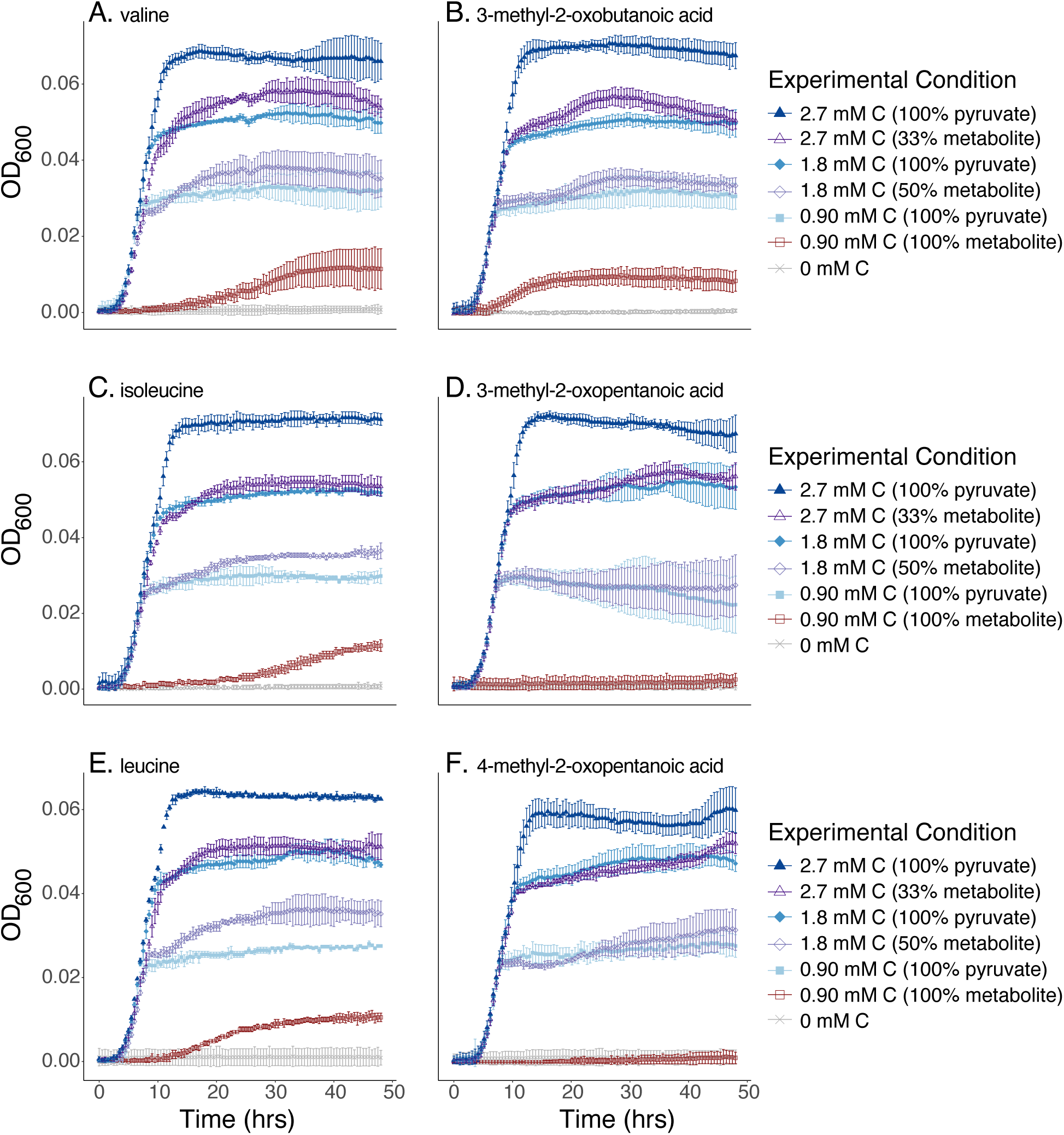
*Alteromonas* growth on a selected *Prochlorococcus*-derived metabolite (red), pyruvate (blue), or a mix of pyruvate and *Prochlorococcus* metabolite (purple). Experiments using only pyruvate are indicated with closed symbols, while experiments using *Prochlorococcus*-derived metabolites are indicated with open symbols. Finally, total moles of added carbon are indicated by squares (0.90 mM C), diamonds (1.8 mM C), or triangles (2.7 mM C). Plot shows mean OD_600_ ± 1 SD (N ≥ 3 except for the leucine, 0.9 mM C condition, where N = 2). Selected metabolites shown here are valine (A); the valine intermediate 3-methyl-2-oxobutanoic acid (B); isoleucine (C); the isoleucine intermediate 3-methyl-2-oxopentanoic acid (D); leucine (E); and the leucine intermediate 4-methyl-2-oxopentanoic acid (F).

**Figure 2.**
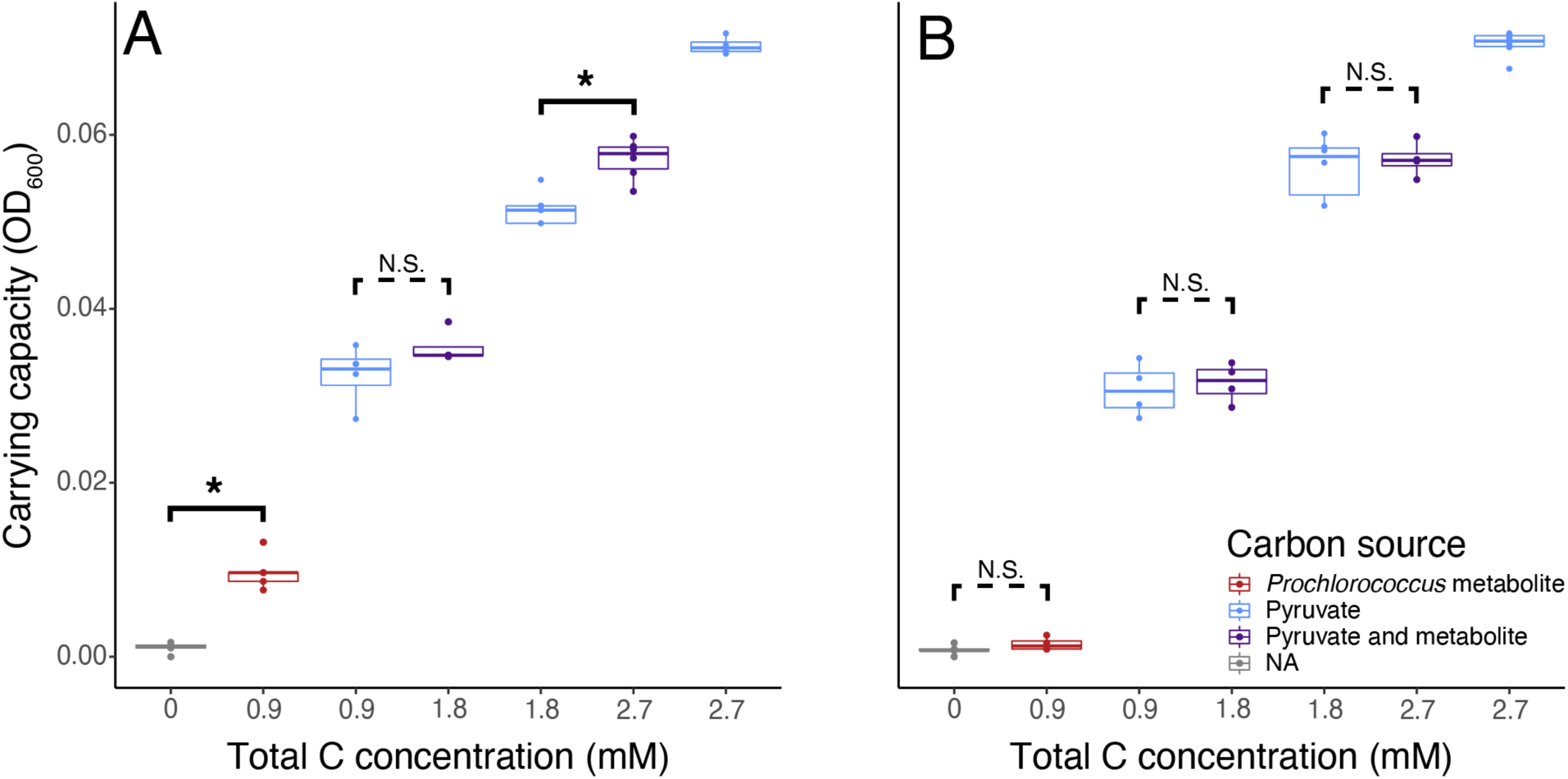
Calculated carrying capacity of *Alteromonas* grown on a selected *Prochlorococcus* metabolite (red), pyruvate (blue), or a mix of pyruvate and the selected metabolite (purple) across a range of carbon concentrations. Example metabolites shown are 3-methyl-2-oxobutanoic acid (A) and 3-methyl-2-oxopentanoic acid (B). The carrying capacity in each carbon condition is significantly different from all other conditions (one way ANOVA with post-hoc Tukey’s HSD, p < 0.05) unless otherwise marked. N.S., not significant. Asterisks indicate a significant difference at the p < 0.05 level.

**Table 1.** Metabolites tested for their ability to support *Alteromonas* growth. *Alteromonas macleodii* MIT1002 was grown with either one of the listed metabolites (all known to be part of *Prochlorococcus* exudates), pyruvate, or a mix of pyruvate and the listed metabolite. A metabolite was considered to support *Alteromonas* growth if carrying capacity was significantly higher when *Alteromonas* cultures received the metabolite than when *Alteromonas* received no *Prochlorococcus* metabolite.

| Metabolite | Supported MIT1002 growth? | Biochemical role |
| --- | --- | --- |
| Leucine | Yes | BCAA metabolism<sup>1</sup> |
| 4-methyl-2-oxopentanoic acid | No | BCAA metabolism |
| Isoleucine | Yes | BCAA metabolism |
| 3-methyl-2-oxopentanoic acid | No | BCAA metabolism |
| Valine | Yes | BCAA metabolism |
| 3-methyl-2-oxobutanoic acid | Yes | BCAA and pantothenate metabolism |
| Kynurenine | No | Tryptophan metabolism |
| Thymidine | No | Pyrimidine metabolism |
| 4-hydroxybenzoic acid | No | Ubiquinone metabolism |

<sup>1</sup>BCAA = branched chain amino acid

### Uptake experiments

To test whether *Alteromonas* is taking up metabolites even when they are not used for net growth, we conducted uptake experiments comparing two structurally and biochemically related metabolites: 3-methyl-2-oxobutanoic acid, the valine precursor which can support *Alteromonas* growth, and 3-methyl-2-oxopentanoic acid, the isoleucine precursor which cannot support *Alteromonas* growth (Figure 1; structures provided in Figure 4). When cultured with 3-methyl-2-oxobutanoic acid, *Alteromonas* cellular abundance increased 94-fold, and 3-methyl-2-oxobutanoic acid concentrations in the media decreased 8-fold over 60 h (Figure 3A). 3-methyl-2-oxobutanoic acid uptake was modeled as a pseudo-first order process after normalizing for cell number, yielding a bulk uptake rate of 0.107 µmol 3-methyl-2-oxobutanoic acid cell^-1^ h^-1^ (Figure S9). In contrast, we observed no significant change in metabolite concentrations in *Alteromonas* cultures incubated with 3-methyl-2-oxopentanoic acid (Figure 3B; Student’s t-test, p > 0.05). These cultures grew slightly at the start of the experiment, presumably from residual seawater carbon, but cell abundance did not increase past the cell abundance observed in controls where no metabolite was added (Figure 3B).

**Figure 3.**
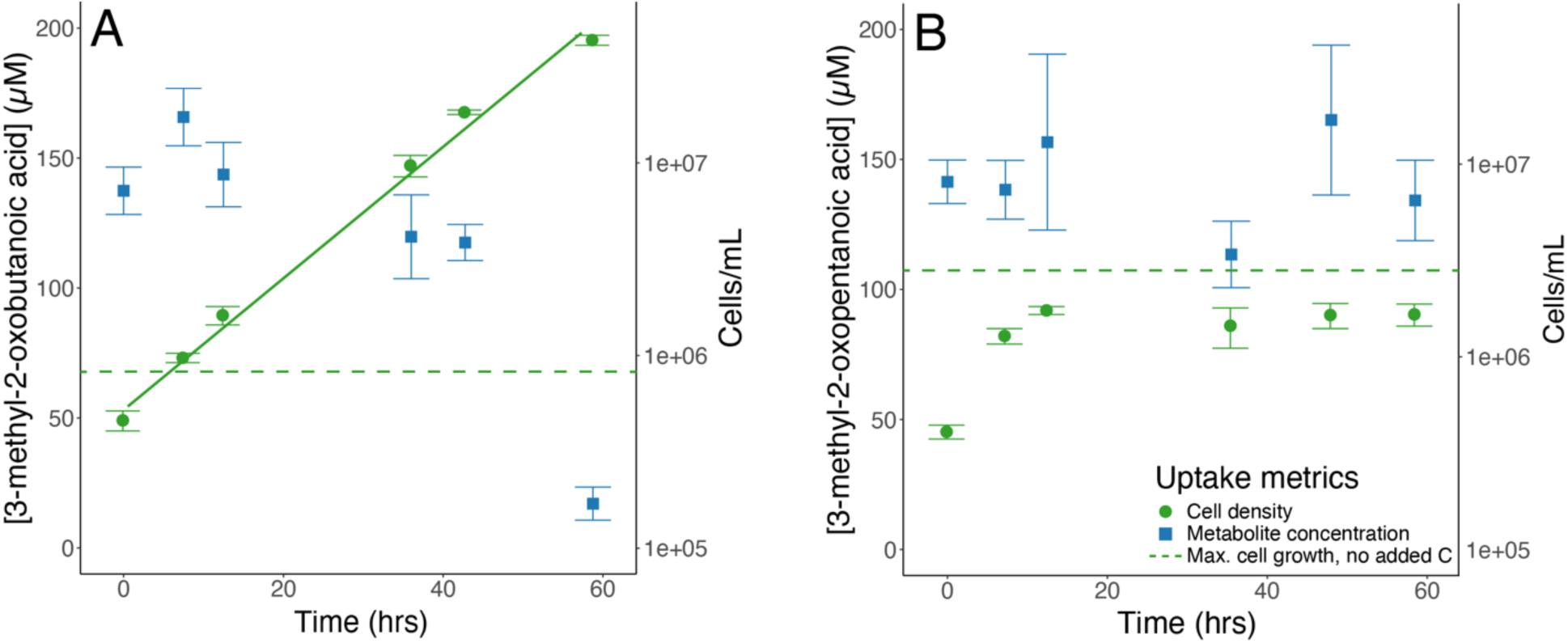
Mean (± 1 SD) *Alteromonas* cell density (green circles) and extracellular metabolite concentration (blue squares) when incubated with either 3-methyl-2-oxobutanoic acid (A) or 3-methyl-2-oxopentanoic acid (B). Dashed green line indicates maximum cell density reached in each experiment by negative controls (i.e., *Alteromonas* cultures incubated in seawater without metabolite addition). Solid line indicates the least squares regression fit to cell density over time for *Alteromonas* cultures grown with 3-methyl-2-oxobutanoic acid (y = 5.74e^0.03x^, where y = cell density and x = time; p < 0.05, R^2^ = 0.99).

**Figure 4.**
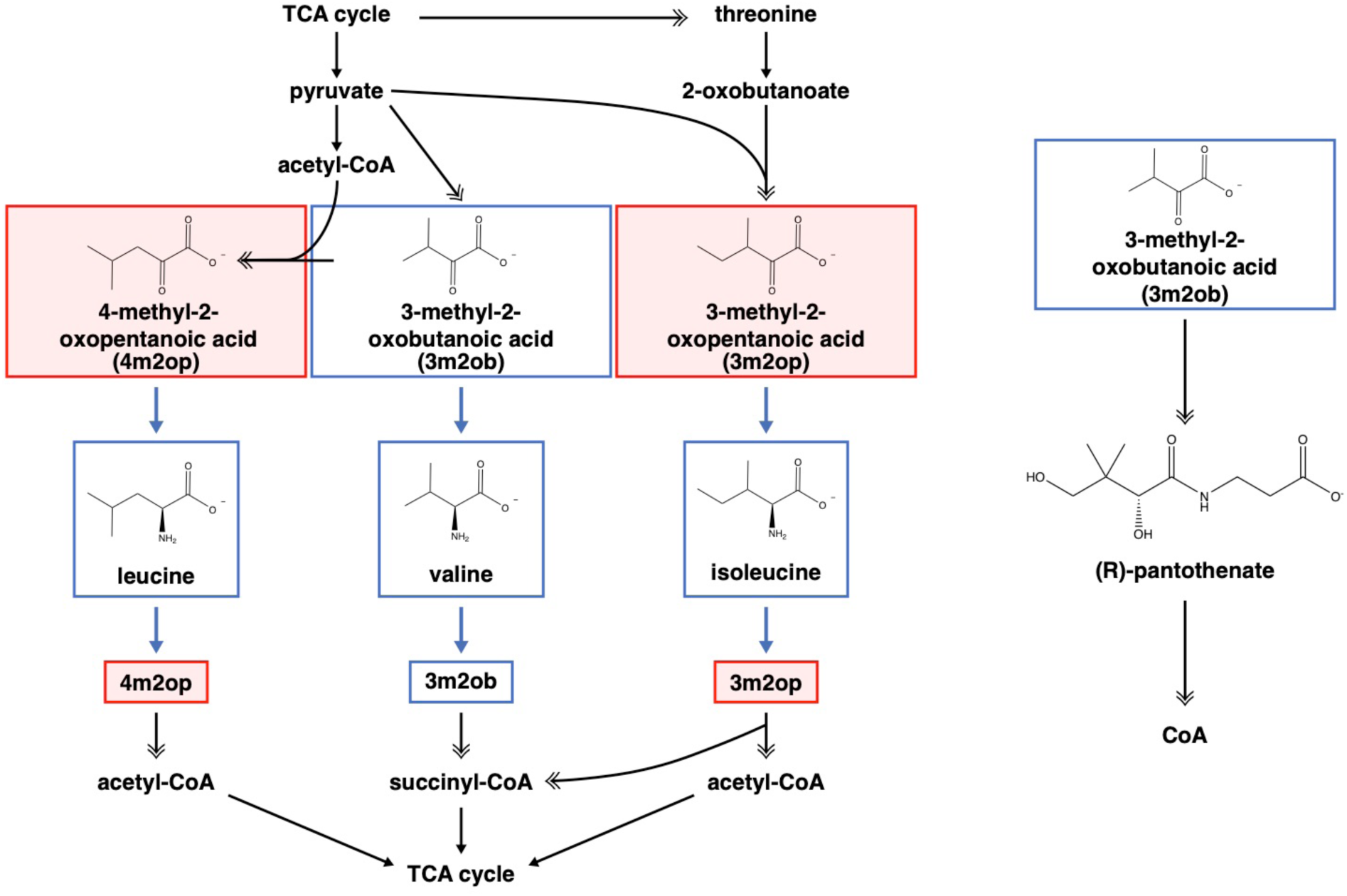
Biosynthetic and catabolic pathways of the branched chain amino acids (left) and of pantothenic acid (right). Metabolites which support *A. macleodii* growth are boxed in blue; metabolites which do not support *A. macleodii* growth are boxed in red. Thick blue arrows indicate a single shared enzyme which mediates transamination between the branched chain amino acids and their immediate precursors/degradation products. Black arrows indicate enzymatic steps, with key intermediates listed; double headed arrows indicate two or more enzymatic reactions. TCA, tricarboxylic acid; CoA, coenzyme A.

## Discussion

While *Alteromonas* is generally recognized to use diverse organic substrates (Noell et al., 2023; Pedler et al., 2014), studies have also highlighted different genetic capabilities and preferences for specific compounds (Koch et al., 2020; Manck et al., 2020; Pedler Sherwood et al., 2015), consistent with our finding that *Alteromonas* only used some of the compounds we tested (Table 1). Still, the specificity of *Alteromonas* response to the branched chain keto acids is notable due to the biochemical and structural similarities of these molecules. *Alteromonas* has the catabolism pathways for all three branched chain amino acids (Figure 4), the first step of which involves a deamination reaction that produces their respective keto acids (Massey et al., 1976; Szentirmai & Horváth, 1976). Further, *Alteromonas* grows on all three branched chain amino acids (Figure 1), indicating it can metabolize all three of the branched chain ketoacids when they are present inside the cell.

In the absence of an intracellular mechanism, transporter specificity likely underlies our observed substrate specificity. The degree of substrate specificity for 3-methyl-2-oxobutanoic acid that we observe in *Alteromonas* is unexpected given previous studies of relevant transporters in marine microbes. A class of 2-oxoacid transporters common in marine microbes can transport the branched chain keto acids, as well as unbranched 2-oxoacids like pyruvate and 2-oxopentanoic acid (Gonin et al., 2007; Pernil et al., 2010; Thomas et al., 2006). Indeed, both ligand binding assays with purified transporters and experiments with transporter mutants have characterized 2-oxoacid transporters which can bind all three of the branched chain keto acids, albeit with slightly higher affinities for 3-methyl-2-oxobutanoic acid than for the other branched chain keto acids (Pernil et al., 2010; Thomas et al., 2006). However, it is not clear which transporter(s) are involved in taking up these compounds in *Alteromonas*, and the promiscuity of transporters can vary widely even within a single family of transporters. For example, one ABC transporter in *Bacillus subtilis* can transport carnitine or choline but not structurally similar osmolytes, while a closely related *B. subtilis* transporter can bind a much wider range of osmolytes (Teichmann et al., 2017). Similarly, five point mutations in a myo-inositol transporter can abolish the transporter affinity for inositol (a 6 carbon sugar) and establish binding for ribose (a 5 carbon sugar) (Herrou & Crosson, 2013). Although to our knowledge a transporter has not been previously reported that can distinguish between 3-methyl-2-oxobutanoic acid and the other branched chain keto acids, *Alteromonas* could feasibly have a transporter that displays this degree of substrate specificity.

Uptake experiments further support our hypothesis of transporter specificity. When cultured with 3-methyl-2-oxobutanoic acid, *Alteromonas* increased in cellular abundance and removed the metabolite at a rate of 0.107 µmol 3-methyl-2-oxobutanoic acid cell^-1^ h^-1^, equivalent to 0.535 µmol C cell^-1^ h^-1^ (Figures 3A, S9). In contrast, *Alteromonas* cultured with 3-methyl-2-oxopentanoic acid did not increase in cellular abundance beyond what was observed in control cultures without a metabolite addition, and we found that *Alteromonas* could not remove 3-methyl-2-oxopentanoic acid from solution (Figure 3B). It is challenging to directly compare these rates to other reported compound-specific and/or microbe-specific uptake rates, due to differences in experimental design (Boysen et al., 2022; Noell & Giovannoni, 2019). However, the 3-methyl-2-oxobutanoic acid uptake rate is in line with the carbon uptake rates observed for *A. macleodii* strain AltSIO when incubated with bulk marine dissolved organic matter (Pedler et al., 2014). This suggests that individual metabolites can recapitulate the biological fluxes of labile bulk DOM, although large differences can exist even between structurally similar metabolites.

The specialization of *Alteromonas* on 3-methyl-2-oxobutanoic acid may reflect a more general role for this metabolite in the microbial ecology of the ocean. 3-methyl-2-oxobutanoic acid was the most abundant branched chain keto acid measured in the exudate of three distinct *Prochlorococcus* strains, and the only branched chain keto acid produced by all three strains (Kujawinski et al., 2023). Additionally, 3-methyl-2-oxobutanoic acid, but not the other branched chain keto acids, has been measured in the dissolved phase in the oligotrophic Sargasso Sea, pointing to a possible broader environmental relevance for this metabolite (Longnecker et al., 2024). This metabolite may be a prevalent substrate for microbial cross-feeding in the ocean, making it potentially advantageous for *Alteromonas* to specialize on 3-methyl-2-oxobutanoic acid.

Biochemically, 3-methyl-2-oxobutanoic acid is also a precursor in pantothenic acid biosynthesis (Figure 4), which further differentiates this metabolite from the other branched chain keto acids. Pantothenic acid is the active component of coenzyme A, which is essential to all life and is used in central carbon metabolism and lipid biosynthesis (Leonardi & Jackowski, 2007). The abundances of both intra- and extracellular pantothenic acid have distinct diel and seasonal patterns in the ocean, suggesting its importance as a marine metabolite (Boysen et al., 2021; Longnecker et al., 2024). The distinctive response of *Alteromonas* to 3-methyl-2-oxobutanoic acid, and the presence of this molecule in the field and in *Prochlorococcus* cultures, could in part reflect its dual role in the metabolism of branched chain amino acids and of pantothenic acid.

For the *Prochlorococcus*-derived metabolites that could support *Alteromonas* growth, we observed diauxic growth curves in cultures which received a mix of pyruvate and metabolite (purple curves, Figure 1A, B, C, E). This biphasic growth pattern suggests that *Alteromonas* uses substrates sequentially, with pyruvate used first. This prioritization effect could be due to the higher growth rate and/or higher carrying capacity when cells were grown on pyruvate relative to other substrates (Figure 1). This is consistent with similar examples of substrate prioritization based on growth rate and/or carrying capacity in other systems (Perrin et al., 2020). The lower carrying capacity from growth on substrates other than pyruvate is seen both in the comparison of cultures grown only on a single substrate and when a portion of carbon from pyruvate was replaced with the other substrates (Figure 2). Differences in carrying capacity between substrates reflects differences in their bacterial growth efficiency (BGE), which is the fraction of carbon in substrates assimilated into biomass. Variations in BGE between substrates has been observed in other systems and can occur as a function of the oxidation state of the substrate, enzymatic steps required for substrate catabolism, energy requirements for substrate transport and metabolism, or genetic capacity of the microbe (Bölscher et al., 2016; Brown & Jones, 2024; del Giorgio & Cole, 1998; Frey et al., 2013; Manzoni et al., 2012; Öquist et al., 2017; Pold et al., 2020; Qiao et al., 2019; Roller & Schmidt, 2015). In the case of *Alteromonas*, we hypothesize that the difference in BGE between substrates is due to differences in enzymatic requirements of associated pathways. That is, while pyruvate can be converted to oxaloacetic acid or acetyl CoA, which feed directly into the citric acid cycle (Voet & Voet, 2011), the catabolism of branched chain amino acids or 3-methyl-2-oxobutanoic acid into central carbon metabolism requires many more enzymatic steps through branched chain coenzyme A intermediates (Massey et al., 1976). This may lead to higher resource or energy investments in using these compounds, potentially lowering the BGE.

## Conclusion

The interactions between microbes and metabolites are crucial to both microbial ecology and carbon cycling. Here we show that *Alteromonas macleodii* MIT1002 grows selectively on *Prochlorococcus*-derived metabolites, likely via transporter specificity. These results further expand our knowledge of the metabolic interactions between *Prochlorococcus* and *Alteromonas*, and offer 3-methyl-2-oxobutanoic acid as a potentially important molecular mediator of these interactions. The specificity of the *Alteromonas* response demonstrates the importance of characterizing microbe-molecule interactions and dissolved organic matter with compound-specific resolution. Small changes in molecular or microbial composition can have an outsized effect on system function, and molecular-level studies enable a nuanced understanding of the larger ocean system that would not be possible through bulk characterization alone.

## Acknowledgements

This work was supported by grants from the Simons Foundation (Award ID #509034 to E.B.K., Award ID #509034SCFY20 to S.W.C. and R.B., SCOPE Award ID #030793 to S.W.C., and Award ID #329108 to M. J. Follows); from the Robert and Ardis James Foundation (to S.W.C); and from the National Science Foundation, via the Center for Chemical Currencies of a Microbial Planet (C-CoMP) Science and Technology Center (NSF award #2019589 to E.B.K). This is C-CoMP publication #053. Support for K.H.H. was provided by a National Defense Science and Engineering Graduate Fellowship. We thank Krista Longnecker, Noah Germolus, Brittany Widner, and Erin McParland for help in lab and helpful conversations; Krista Longnecker, Alex Frank, and Helen Fredricks for flow cytometry expertise; and Ben Van Mooy for use of his flow cytometer.

## Notes

https://github.com/khhalloran/Alteromonas_mtab_uptake

### Competing Interest Statement

The authors have declared no competing interest.

## Works Cited

Aharonovich, D., & Sher, D. (2016). Transcriptional response of *Prochlorococcus* to co-culture with a marine *Alteromonas*: Differences between strains and the involvement of putative infochemicals. The ISME Journal, 10(12), 2892–2906. 10.1038/ismej.2016.70

Becker, J. W., Hogle, S. L., Rosendo, K., & Chisholm, S. W. (2019). Co-culture and biogeography of *Prochlorococcus* and SAR11. The ISME Journal, 13(6), 1506–1519. 10.1038/s41396-019-0365-4

Bertilsson, S., Berglund, O., Pullin, M. J., & Chisholm, S. W. (2005). Release of dissolved organic matter by *Prochlorococcus*. Vie et Milieu/Life & Environment, 225–231.

Berube, P. M., Biller, S. J., Kent, A. G., Berta-Thompson, J. W., Roggensack, S. E., Roache-Johnson, K. H., Ackerman, M., Moore, L. R., Meisel, J. D., Sher, D., Thompson, L. R., Campbell, L., Martiny, A. C., & Chisholm, S. W. (2015). Physiology and evolution of nitrate acquisition in *Prochlorococcus*. The ISME Journal, 9(5), 1195–1207. 10.1038/ismej.2014.211

Biller, S. J., Coe, A., & Chisholm, S. W. (2016). Torn apart and reunited: Impact of a heterotroph on the transcriptome of *Prochlorococcus*. The ISME Journal, 10(12), 2831–2843. 10.1038/ismej.2016.82

Biller, S. J., Coe, A., Martin-Cuadrado, A.-B., & Chisholm, S. W. (2015). Draft genome sequence of *Alteromonas macleodii* strain MIT1002, isolated from an enrichment culture of the marine cyanobacterium *Prochlorococcus*. Genome Announcements, 3(4), e00967–15, /ga/3/4/e00967-15.atom. 10.1128/genomeA.00967-15

Bölscher, T., Wadsö, L., Börjesson, G., & Herrmann, A. M. (2016). Differences in substrate use efficiency: Impacts of microbial community composition, land use management, and substrate complexity. Biology and Fertility of Soils, 52(4), 547–559. 10.1007/s00374-016-1097-5

Boysen, A. K., Carlson, L. T., Durham, B. P., Groussman, R. D., Aylward, F. O., Ribalet, F., Heal, K. R., White, A. E., DeLong, E. F., Armbrust, E. V., & Ingalls, A. E. (2021). Particulate metabolites and transcripts reflect diel oscillations of microbial activity in the surface ocean. mSystems, 6(3), e00896–20. 10.1128/mSystems.00896-20

Boysen, A. K., Durham, B. P., Kumler, W., Key, R. S., Heal, K. R., Carlson, L. T., Groussman, R. D., Armbrust, E. V., & Ingalls, A. E. (2022). Glycine betaine uptake and metabolism in marine microbial communities. Environmental Microbiology, 24(5), 2380–2403. 10.1111/1462-2920.16020

Brown, R. W., & Jones, D. L. (2024). Plasticity of microbial substrate carbon use efficiency in response to changes in plant carbon input and soil organic matter status. Soil Biology and Biochemistry, 188, 109230. 10.1016/j.soilbio.2023.109230

Calfee, B. C., Glasgo, L. D., & Zinser, E. R. (2022). *Prochlorococcus* exudate stimulates heterotrophic bacterial competition with rival phytoplankton for available nitrogen. mBio, 13(1), e02571–21. 10.1128/mbio.02571-21

del Giorgio, P. A., & Cole, J. J. (1998). Bacterial growth efficiency in natural aquatic systems. Annual Review of Ecology and Systematics, 29(1), 503–541. 10.1146/annurev.ecolsys.29.1.503

Dittmar, T., Koch, B., Hertkorn, N., & Kattner, G. (2008). A simple and efficient method for the solid-phase extraction of dissolved organic matter (SPE-DOM) from seawater. Limnology and Oceanography: Methods, 6(6), 230–235. 10.4319/lom.2008.6.230

Flombaum, P., Gallegos, J. L., Gordillo, R. A., Rincón, J., Zabala, L. L., Jiao, N., Karl, D. M., Li, W. K. W., Lomas, M. W., Veneziano, D., Vera, C. S., Vrugt, J. A., & Martiny, A. C. (2013). Present and future global distributions of the marine cyanobacteria *Prochlorococcus* and *Synechococcus*. Proceedings of the National Academy of Sciences, 110(24), 9824–9829. 10.1073/pnas.1307701110

Forchielli, E., Sher, D., & Segrè, D. (2022). Metabolic phenotyping of marine heterotrophs on refactored media reveals diverse metabolic adaptations and lifestyle strategies. mSystems, 7(4), e00070–22. 10.1128/msystems.00070-22

Frey, S. D., Lee, J., Melillo, J. M., & Six, J. (2013). The temperature response of soil microbial efficiency and its feedback to climate. Nature Climate Change, 3(4), 395–398. 10.1038/nclimate1796

Givati, S., Forchielli, E., Aharonovich, D., Barak, N., Weissberg, O., Belkin, N., Rahav, E., Segrè, D., & Sher, D. (2024). Diversity in the utilization of different molecular classes of dissolved organic matter by heterotrophic marine bacteria. Applied and Environmental Microbiology, 90(7), e00256–24. 10.1128/aem.00256-24

Gonin, S., Arnoux, P., Pierru, B., Lavergne, J., Alonso, B., Sabaty, M., & Pignol, D. (2007). Crystal structures of an extracytoplasmic solute receptor from a TRAP transporter in its open and closed forms reveal a helix-swapped dimer requiring a cation for α-keto acid binding. BMC Structural Biology, 7(1), 11. 10.1186/1472-6807-7-11

Hansell, D., Carlson, C., Repeta, D., & Schlitzer, R. (2009). Dissolved organic matter in the ocean: A controversy stimulates new insights. Oceanography, 22(4), 202–211. 10.5670/oceanog.2009.109

Herrou, J., & Crosson, S. (2013). *Myo*-inositol and D-ribose ligand discrimination in an ABC periplasmic binding protein. Journal of Bacteriology, 195(10), 2379–2388. 10.1128/JB.00116-13

Ivars-Martinez, E., Martin-Cuadrado, A.-B., D’Auria, G., Mira, A., Ferriera, S., Johnson, J., Friedman, R., & Rodriguez-Valera, F. (2008). Comparative genomics of two ecotypes of the marine planktonic copiotroph *Alteromonas macleodii* suggests alternative lifestyles associated with different kinds of particulate organic matter. The ISME Journal, 2(12), 1194–1212. 10.1038/ismej.2008.74

Kido Soule, M. C., Longnecker, K., Johnson, W. M., & Kujawinski, E. B. (2015). Environmental metabolomics: Analytical strategies. Marine Chemistry, 177, 374–387. 10.1016/j.marchem.2015.06.029

Koch, H., Germscheid, N., Freese, H. M., Noriega-Ortega, B., Lücking, D., Berger, M., Qiu, G., Marzinelli, E. M., Campbell, A. H., Steinberg, P. D., Overmann, J., Dittmar, T., Simon, M., & Wietz, M. (2020). Genomic, metabolic and phenotypic variability shapes ecological differentiation and intraspecies interactions of *Alteromonas macleodii*. Scientific Reports, 10(1), 809. 10.1038/s41598-020-57526-5

Kujawinski, E. B., Braakman, R., Longnecker, K., Becker, J. W., Chisholm, S. W., Dooley, K., Kido Soule, M. C., Swarr, G. J., & Halloran, K. (2023). Metabolite diversity among representatives of divergent *Prochlorococcus* ecotypes. mSystems, 8(5), e01261–22. 10.1128/msystems.01261-22

Leonardi, R., & Jackowski, S. (2007). Biosynthesis of pantothenic acid and Coenzyme A. EcoSal Plus, 2(2), ecosalplus.3.6.3.4. 10.1128/ecosalplus.3.6.3.4

Letscher, R. T., Moore, J. K., Martiny, A. C., & Lomas, M. W. (2023). Biodiversity and stoichiometric plasticity increase pico-phytoplankton contributions to marine net primary productivity and the biological pump. Global Biogeochemical Cycles, 37(8), e2023GB007756. 10.1029/2023GB007756

Liu, S., Baetge, N., Comstock, J., Opalk, K., Parsons, R., Halewood, E., English, C. J., Giovannoni, S., Bolaños, L. M., Nelson, C. E., Vergin, K., & Carlson, C. A. (2020). Stable isotope probing identifies bacterioplankton lineages capable of utilizing dissolved organic matter across a range of bioavailability. Frontiers in Microbiology, 11, 580397. 10.3389/fmicb.2020.580397

Longnecker, K., Kido Soule, M. C., Swarr, G. J., Parsons, R. J., Liu, S., Johnson, W. M., Widner, B., Curry, R., Carlson, C. A., & Kujawinski, E. B. (2024). Seasonal and daily patterns in known dissolved metabolites in the northwestern Sargasso Sea. Limnology and Oceanography, 69(3), 449–466. 10.1002/lno.12497

López-Sandoval, D., Rodríguez-Ramos, T., Cermeño, P., & Marañón, E. (2013). Exudation of organic carbon by marine phytoplankton: Dependence on taxon and cell size. Marine Ecology Progress Series, 477, 53–60. 10.3354/meps10174

Manck, L.E., Espinoza, J.L., Dupont, C.L., and Barbeau, K.A. (2020). Transcriptomic study of substrate-specific transport mechanisms for iron and carbon in the marine copiotroph Alteromonas macleodii. mSystems, 5(2), e00070–20. 10.1128/mSystems.00070-20

Manzoni, S., Taylor, P., Richter, A., Porporato, A., & Ågren, G. I. (2012). Environmental and stoichiometric controls on microbial carbon-use efficiency in soils. New Phytologist, 196(1), 79–91. 10.1111/j.1469-8137.2012.04225.x

Massey, L. K., Sokatch, J. R., & Conrad, R. S. (1976). Branched-chain amino acid catabolism in bacteria. Bacteriological Reviews, 40(1), 42–54.

Moran, M. A., Ferrer-González, F. X., Fu, H., Nowinski, B., Olofsson, M., Powers, M. A., Schreier, J. E., Schroer, W. F., Smith, C. B., & Uchimiya, M. (2022). The ocean’s labile DOC supply chain. Limnology and Oceanography, 67(5), 1007–1021. 10.1002/lno.12053

Moran, M. A., Kujawinski, E. B., Schroer, W. F., Amin, S. A., Bates, N. R., Bertrand, E. M., Braakman, R., Brown, C. T., Covert, M. W., Doney, S. C., Dyhrman, S. T., Edison, A. S., Eren, A. M., Levine, N. M., Li, L., Ross, A. C., Saito, M. A., Santoro, A. E., Segrè, D.,… Vardi, A. (2022). Microbial metabolites in the marine carbon cycle. Nature Microbiology, 7(4), 508–523. 10.1038/s41564-022-01090-3

Morris, J. J., Johnson, Z. I., Szul, M. J., Keller, M., & Zinser, E. R. (2011). Dependence of the cyanobacterium *Prochlorococcus* on hydrogen peroxide scavenging microbes for growth at the ocean’s surface. PLoS ONE, 6(2), e16805. 10.1371/journal.pone.0016805

Morris, J. J., Kirkegaard, R., Szul, M. J., Johnson, Z. I., & Zinser, E. R. (2008). Facilitation of robust growth of *Prochlorococcus* colonies and dilute liquid cultures by “helper” heterotrophic bacteria. Applied and Environmental Microbiology, 74(14), 4530–4534. 10.1128/AEM.02479-07

Morris, J. J., Lenski, R. E., & Zinser, E. R. (2012). The black queen hypothesis: Evolution of dependencies through adaptive gene loss. mBio, 3(2), e00036–12. 10.1128/mBio.00036-12

Noell, S. E., Brennan, E., Washburn, Q., Davis, E. W., Hellweger, F. L., & Giovannoni, S. J. (2023). Differences in the regulatory strategies of marine oligotrophs and copiotrophs reflect differences in motility. Environmental Microbiology, 25(7), 1265–1280. 10.1111/1462-2920.16357

Noell, S. E., & Giovannoni, S. J. (2019). SAR11 bacteria have a high affinity and multifunctional glycine betaine transporter. Environmental Microbiology, 21(7), 2559– 2575. 10.1111/1462-2920.14649

Ofaim, S., Sulheim, S., Almaas, E., Sher, D., & Segrè, D. (2021). Dynamic allocation of carbon storage and nutrient-dependent exudation in a revised genome-scale model of *Prochlorococcus*. Frontiers in Genetics, 12, 586293. 10.3389/fgene.2021.586293

Öquist, M. G., Erhagen, B., Haei, M., Sparrman, T., Ilstedt, U., Schleucher, J., & Nilsson, M. B. (2017). The effect of temperature and substrate quality on the carbon use efficiency of saprotrophic decomposition. Plant and Soil, 414(1–2), 113–125. 10.1007/s11104-016-3104-x

Partensky, F., Hess, W. R., & Vaulot, D. (1999). *Prochlorococcus*, a marine photosynthetic prokaryote of global significance. Microbiology and Molecular Biology Reviews, 63(1), 106.

Pedler, B. E., Aluwihare, L. I., & Azam, F. (2014). Single bacterial strain capable of significant contribution to carbon cycling in the surface ocean. Proceedings of the National Academy of Sciences, 111(20), 7202–7207. 10.1073/pnas.1401887111

Pedler Sherwood, B., Shaffer, E. A., Reyes, K., Longnecker, K., Aluwihare, L. I., & Azam, F. (2015). Metabolic characterization of a model heterotrophic bacterium capable of significant chemical alteration of marine dissolved organic matter. Marine Chemistry, 177, 357–365. 10.1016/j.marchem.2015.06.027

Pernil, R., Herrero, A., & Flores, E. (2010). A TRAP transporter for pyruvate and other monocarboxylate 2-oxoacids in the cyanobacterium *Anabaena* sp. strain PCC 7120. Journal of Bacteriology, 192(22), 6089–6092. 10.1128/JB.00982-10

Perrin, E., Ghini, V., Giovannini, M., Di Patti, F., Cardazzo, B., Carraro, L., Fagorzi, C., Turano, P., Fani, R., & Fondi, M. (2020). Diauxie and co-utilization of carbon sources can coexist during bacterial growth in nutritionally complex environments. Nature Communications, 11(1), 3135. 10.1038/s41467-020-16872-8

Pold, G., Domeignoz-Horta, L. A., Morrison, E. W., Frey, S. D., Sistla, S. A., & DeAngelis, K. M. (2020). Carbon use efficiency and its temperature sensitivity covary in soil bacteria. mBio, 11(1), e02293–19. 10.1128/mBio.02293-19

Qiao, Y., Wang, J., Liang, G., Du, Z., Zhou, J., Zhu, C., Huang, K., Zhou, X., Luo, Y., Yan, L., & Xia, J. (2019). Global variation of soil microbial carbon-use efficiency in relation to growth temperature and substrate supply. Scientific Reports, 9(1), 5621. 10.1038/s41598-019-42145-6

Roller, B. R. K., & Schmidt, T. M. (2015). The physiology and ecological implications of efficient growth. The ISME Journal, 9(7), 1481–1487. 10.1038/ismej.2014.235

Saifuddin, M., Bhatnagar, J. M., Segrè, D., & Finzi, A. C. (2019). Microbial carbon use efficiency predicted from genome-scale metabolic models. Nature Communications, 10(1), 3568. 10.1038/s41467-019-11488-z

Sher, D., Thompson, J. W., Kashtan, N., Croal, L., & Chisholm, S. W. (2011). Response of *Prochlorococcus* ecotypes to co-culture with diverse marine bacteria. The ISME Journal, 5(7), 1125–1132. 10.1038/ismej.2011.1

Szentirmai, A., & Horváth, I. (1976). Regulation of branched-chain amino acid biosynthesis. Acta Microbiologica Academiae Scientiarum Hungaricae, 23(2), 137–149.

Szul, M. J., Dearth, S. P., Campagna, S. R., & Zinser, E. R. (2019). Carbon fate and flux in *Prochlorococcus* under nitrogen limitation. mSystems, 4(1), e00254–18. 10.1128/mSystems.00254-18

Teichmann, L., Chen, C., Hoffmann, T., Smits, S. H. J., Schmitt, L., & Bremer, E. (2017). From substrate specificity to promiscuity: Hybrid ABC transporters for osmoprotectants. Molecular Microbiology, 104(5), 761–780. 10.1111/mmi.13660

Thomas, G. H., Southworth, T., León-Kempis, M. R., Leech, A., & Kelly, D. J. (2006). Novel ligands for the extracellular solute receptors of two bacterial TRAP transporters. Microbiology, 152(1), 187–198. 10.1099/mic.0.28334-0

Voet, D., & Voet, J. G. (2011). Biochemistry (4. ed). Wiley.

