## Supplemental methods; Supplemental Figures 1-9 for "Uptake of Prochlorococcus-derived metabolites by *Alteromonas macleodii* MIT1002 shows high levels of substrate specificity"

### Supplemental Information

#### Supplemental Methods

##### Materials

Sodium acetate (≥ 99%), sodium lactate (≥ 99%), sodium pyruvate (≥ 99%), glycerol (≥ 99%), thymidine (≥ 99%), 3-methyl-2-oxobutanoic acid (95%), 4-hydroxybenzoic acid (≥ 99%), kynurenine (≥ 98%), 3-methyl-2-oxopentanoic acid (≥ 98%), 4-methyl-2-oxopentanoic acid (≥ 98%), leucine (≥ 99.5%), isoleucine (≥ 99.5%), valine (≥ 99.5%), ammonium chloride (> 99.5%), sodium phosphate (≥ 99.5%), EDTA (≥ 99%), and iron (III) chloride (≥ 98%) were purchased from Sigma-Aldrich (St. Louis, MO, USA). Deionized water was obtained from a Milli-Q system (Millipore; resistivity 18.2 MΩ at 24˚C, TOC < 1 µM). Organic substrate and nutrient stocks were filtered through 0.2 µm nylon syringe filters (Fisher or Corning) under sterile conditions prior to addition to culture media. Cultures were maintained and uptake experiments were performed in 125 mL polycarbonate bottles, which were acid washed in 10% hydrochloric acid and autoclaved before use.

##### Growth Experiment Optimization

Prior to uptake experiments, experiments were performed to confirm *Alteromonas* growth on minimal media and lower-carbon media, and to determine the change in carbon concentration required to detect a statistically significant difference in culture carrying capacity. All optimization experiments contained the same inorganic nutrients as found in ProMM (Berube et al., 2015). Experiments were performed on black walled 96-well plates (Corning, Arizona) using a BioTek Synergy 2 plate reader, measuring $OD_{600}$ every 30 minutes for 48 hours.

To test the ability of *Alteromonas* to grow on minimal media, we grew cultures under one of three conditions: ProMM (which contains pyruvate, acetate, lactate, and glycerol), pyruvate with vitamins, or pyruvate alone. For the "pyruvate" and "pyruvate with vitamins" conditions, pyruvate was added to 6.9 mM carbon, matching the total carbon concentration in ProMM. For the "pyruvate and vitamins" condition, vitamins were added at the same concentration as ProMM. Over the course of 48 hours, there was no significant difference in culture carrying capacity between conditions (ANOVA, $p > 0.05$, Figure S1).

To test the effect of growing *Alteromonas* on lower carbon concentrations than found in ProMM, we grew cultured on either 6.8, 5.5, 5, 4.1, 2.7, or 0 mM carbon from pyruvate. We found that *Alteromonas* could grow on lower carbon concentrations than those in ProMM (Figure S2), and that changing carbon concentrations by 0.9 mM C was sufficient for observing a significant difference in carrying capacity.

**Flow Cytometry**

Samples for measuring cell number by flow cytometry were stained for 10 minutes with SYBR Green 1 (1X final concentration) and diluted with a Tris-HCl/EDTA buffer to approximately $5x10^4$ cells $mL^{-1}$ prior to analysis. Cells were then quantified using a Guava easyCyte HT flow cytometer (Millipore). Instrument flow rate was set to medium ($0.59\ \mu L\ s^{-1}$). *Alteromonas* was gated using green fluorescence and side scatter using an elliptical gate that encompassed the range of the population. All flow cytometry files were analyzed using guavaSoft InCyte 2.7.

**Liquid Chromatography-tandem mass spectrometry (LC-MS/MS)**

Samples were analyzed in randomized order by ultra-performance liquid chromatography (UPLC, Accela Open Autosampler and Accela 1250 Pump, Thermo Scientific) coupled via heated electrospray ionization (H-ESI) to a triple quadrupole mass spectrometer (TSQ Vantage, Thermo Scientific) operated under selected reaction monitoring (SRM) mode. Spray voltage was set at 4000 V (positive mode) and 3200 V (negative mode). Source gases were set at 55 (sheath) and 20 (auxiliary), while the heated capillary temperature was 375 °C, and the vaporizer temperature was 400 °C. Chromatographic separation was performed on a Waters Acquity HSS T3 column (2.1 × 100 mm, 1.8 μm) equipped with a Vanguard pre-column and maintained at 40 °C. The column was eluted with (A) 0.1% formic acid in water and (B) 0.1% formic acid in acetonitrile at a flow rate of 0.5 mL $min^{-1}$.The gradient starts at 1% B for 1 min, ramps to 15% B from 1-3 min, ramps to 50% from 3-6 min, ramps to 95% B from 6-9 min, holds at 95% B until 10 min, ramps to 1% from 10-10.2 min, with re-equilibration at 1% B (total gradient time 12 min). Separate autosampler injections of 5 μL each were made for positive and negative ion modes. Pooled samples were run every five samples to act as a quality control. Raw files were converted to mzML using MSconvert (Chambers et al., 2012), peak areas were integrated in EL-MAVEN (Agrawal et al., 2019), curve fitting and quantification was performed in MATLAB, and statistical analysis was performed in R (R Core Team, 2021).

#### Supplemental Figures

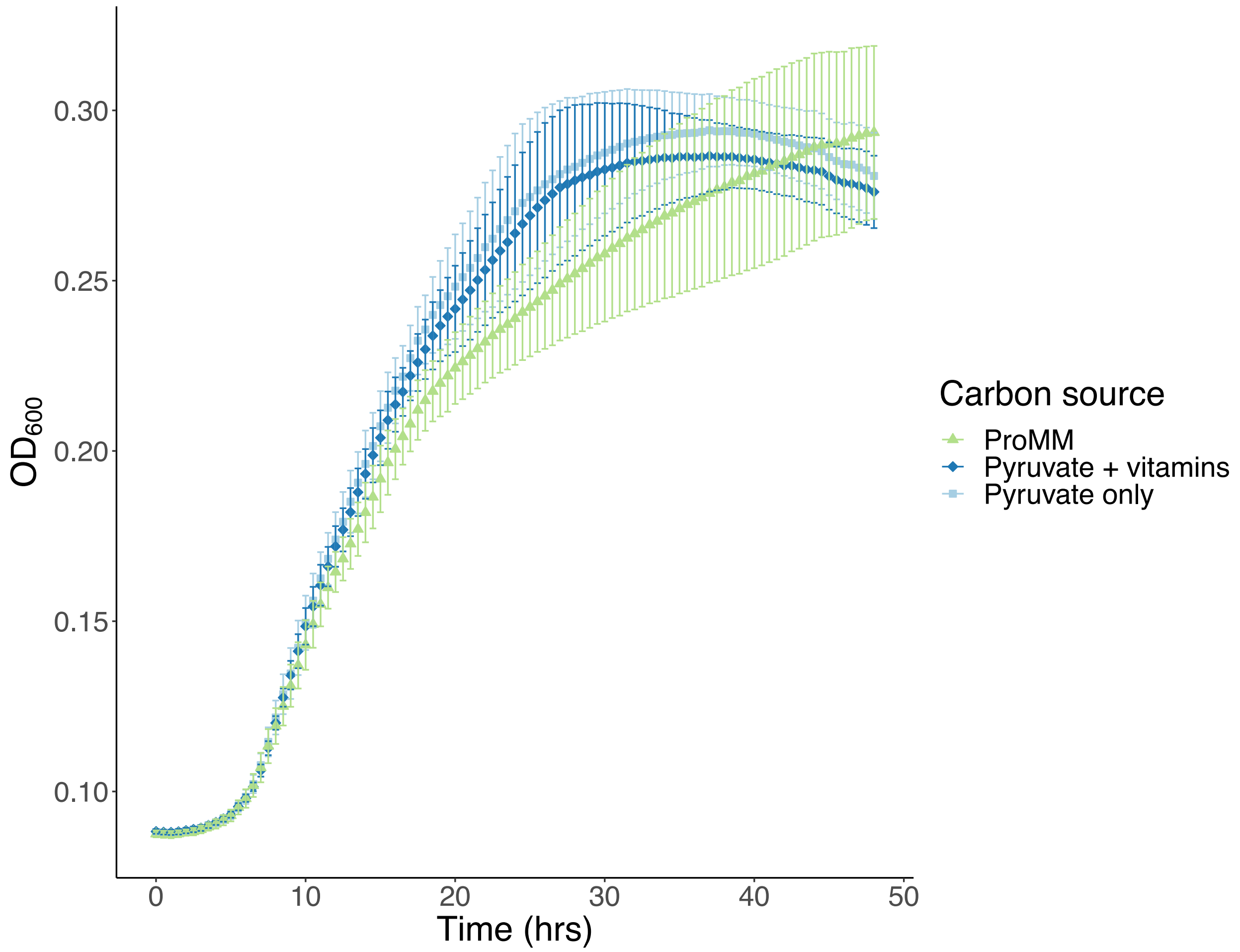


Figure S1. *Alteromonas* growth over 48 hours on either ProMM media, pyruvate (6.9 mM carbon, matching total ProMM carbon concentration) and vitamins (Va vitamin mix, matching the ProMM vitamin addition), or pyruvate alone (6.9 mM carbon). Carrying capacity was not significantly different between conditions (ANOVA, $p > 0.05$).

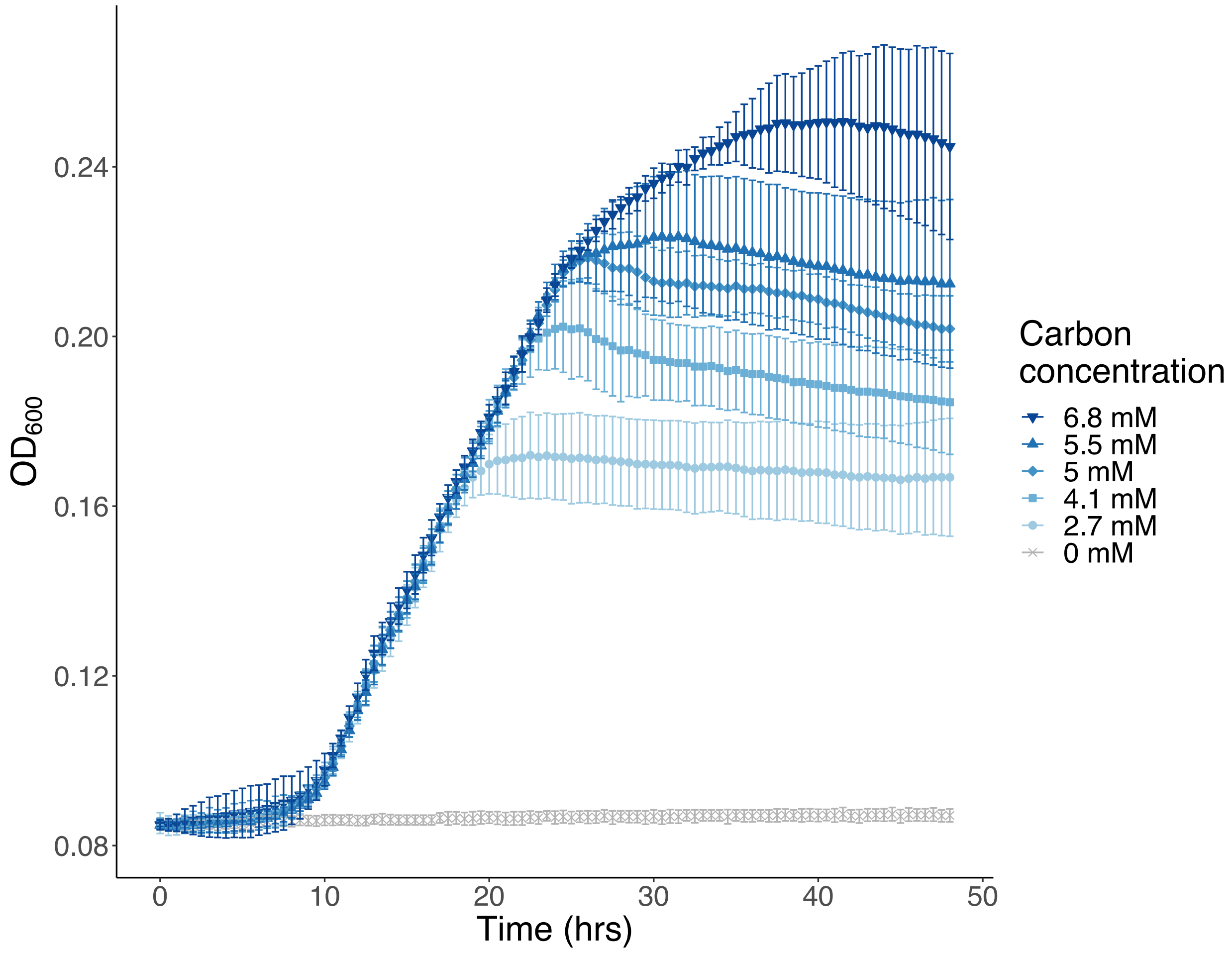


Figure S2. *Alteromonas* growth on varying concentrations of carbon from pyruvate, from 6.8 mM to 2.7 mM carbon.

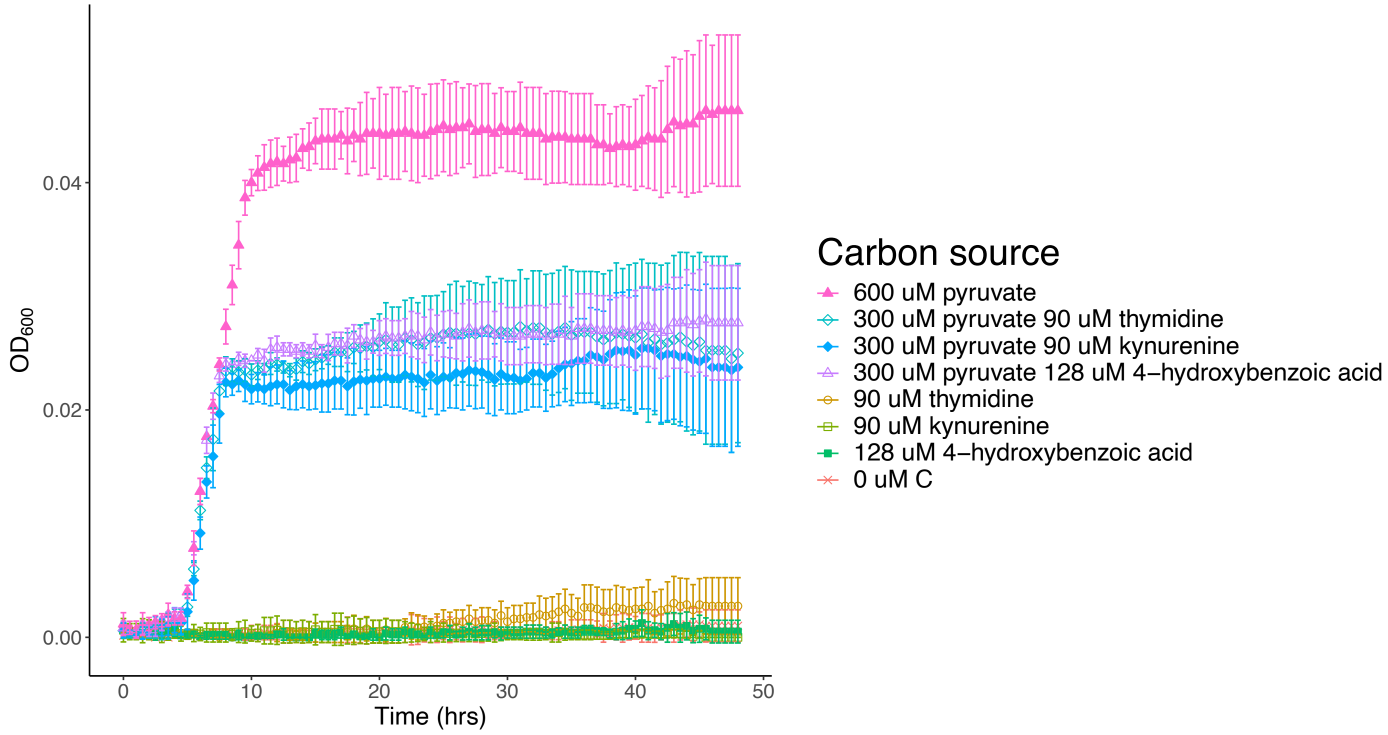


Figure S3. Growth curves of *Alteromonas* incubated with metabolites which cannot support growth. *Alteromonas* was grown with either 600 µM pyruvate as a positive control, no added carbon as a negative control, an example metabolite (4-hydroxybenzoic acid, kynurenine, or thymidine) which were could not support growth in initial experiments, or a mix of pyruvate and an example metabolite. Total carbon concentrations in the mixed substrate cultures are equivalent to the carbon received from 600 µM pyruvate (1.8 mM).

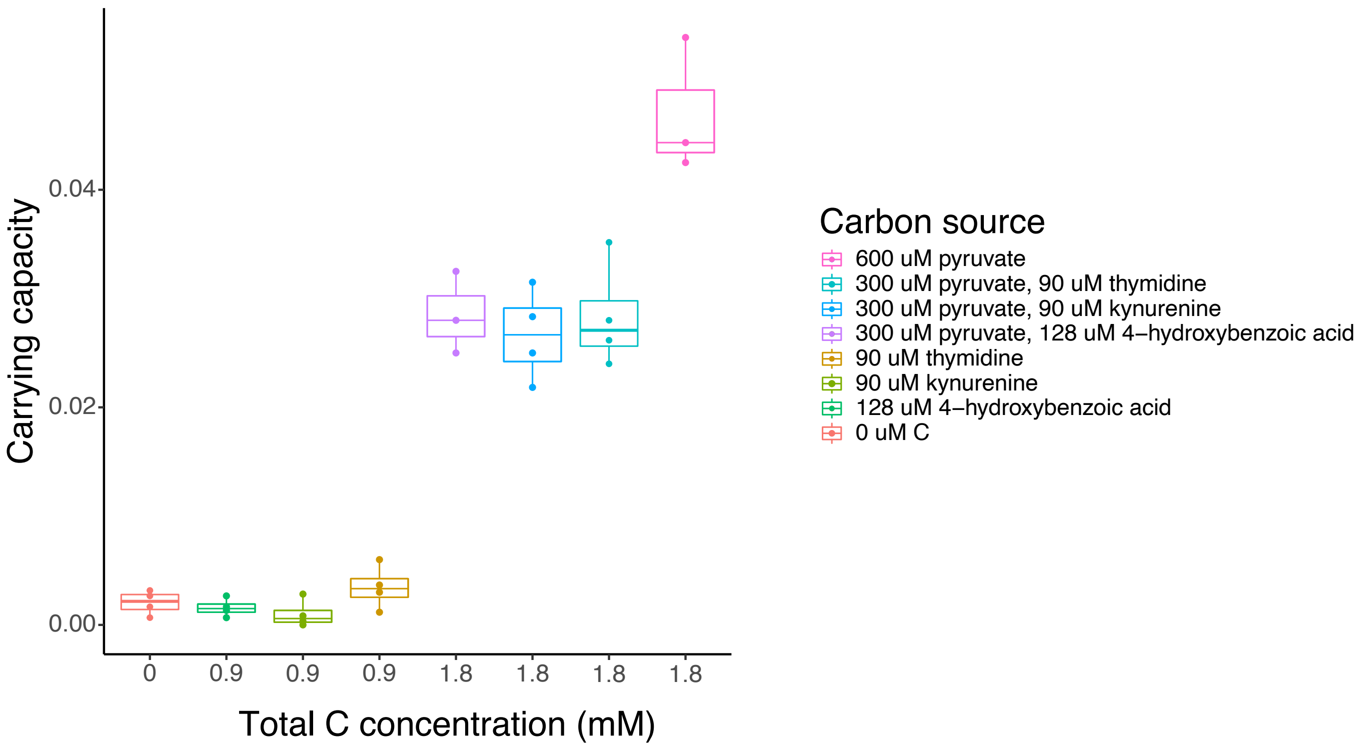


Figure S4. Calculated carrying capacities of *Alteromonas* cultures incubated with pyruvate, a *Prochlorococcus* metabolite which cannot support growth, or a mix of pyruvate and one of these metabolites. There was no significant difference in carrying capacity between cultures which received 0 mM carbon and those which received 0.9 mM C from any of the three carbon sources, and no significant difference in carrying capacity between cultures which received 1.8 mM carbon from a mix of pyruvate and any one metabolite (two way ANOVA with post-hoc Tukey's HSD test, $p > 0.05$). *Alteromonas* grown on 1.8 mM C from pyruvate reached significantly higher carrying capacities than *Alteromonas* growth with 1.8 mM C from a mix of pyruvate and a *Prochlorococcus* metabolite (two way ANOVA with post-hoc Tukey's HSD test, $p < 0.05$).

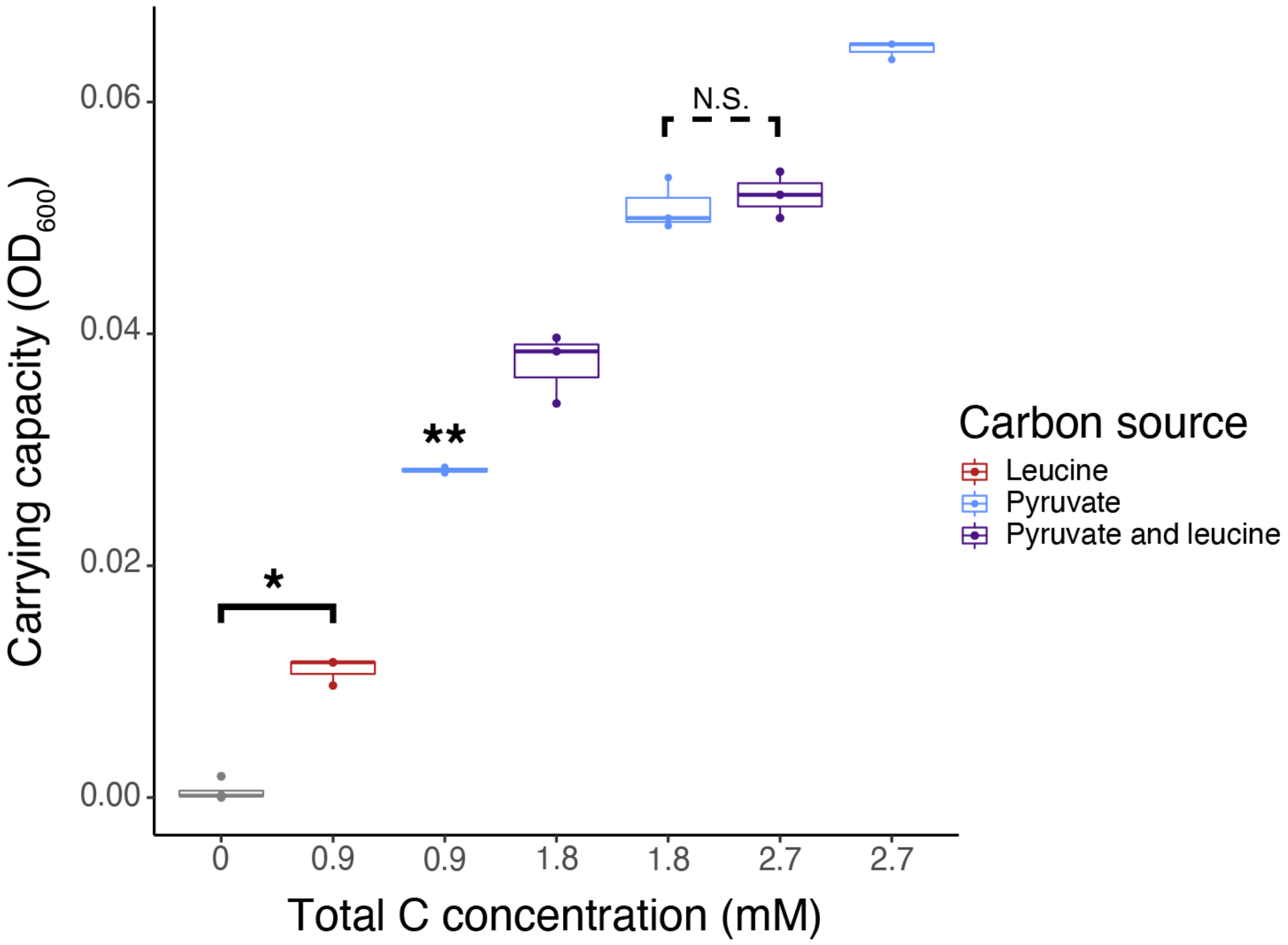


Figure S5. Calculated carrying capacities of *Alteromonas* incubated with leucine (red), pyruvate (blue), or a mix of pyruvate and leucine (purple). All carrying capacities are significantly different from each other unless otherwise marked (two way ANOVA with post-hoc Tukey's HSD, $p < 0.05$). N.S., not significant. Asterisk indicates a significant difference at the $p < 0.05$ level. Double asterisk indicates a condition with only 2 replicates, due to removal of wells from analysis while accounting for growth plate position effects.

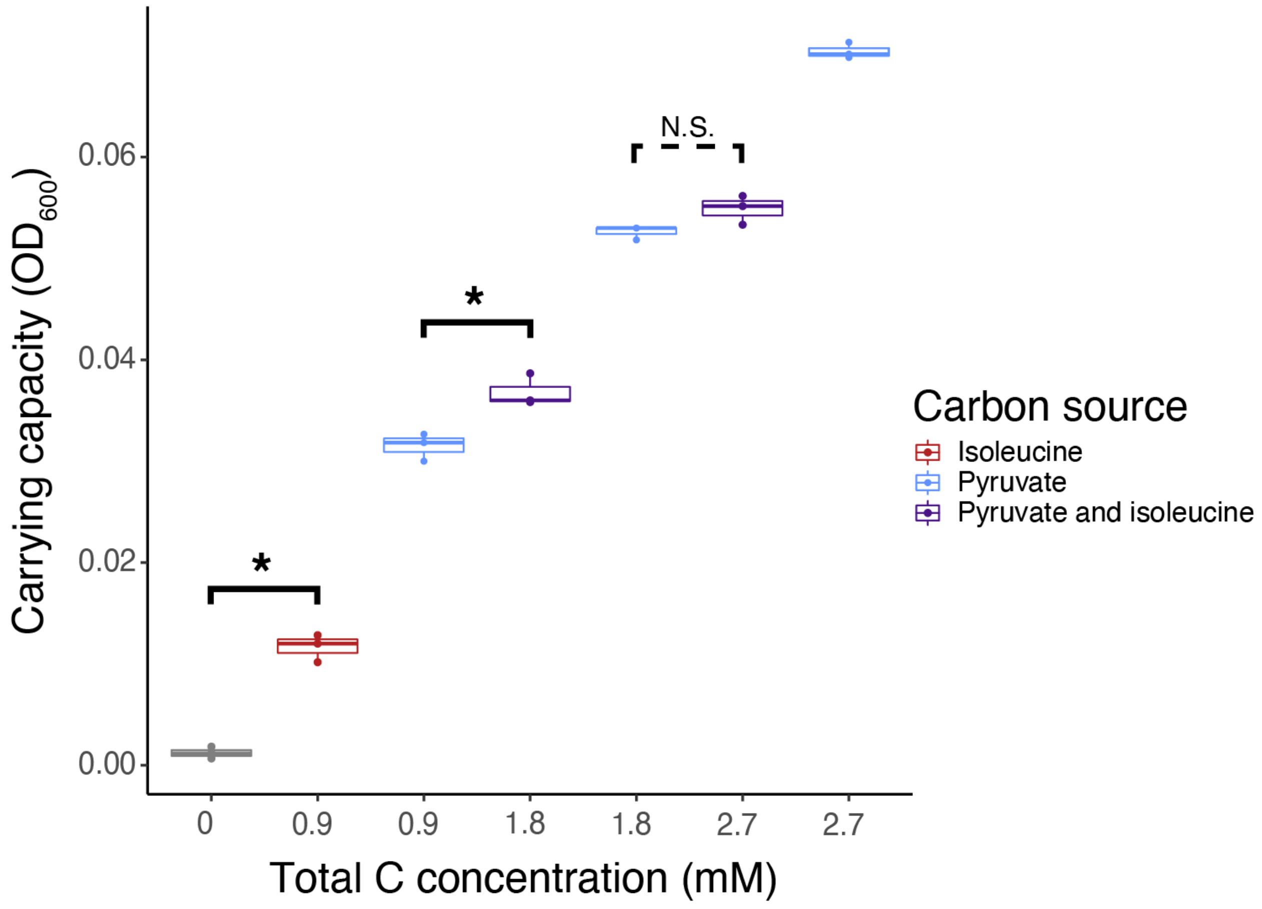


Figure S6. Calculated carrying capacities of *Alteromonas* incubated with isoleucine (red), pyruvate (blue), or a mix of pyruvate and isoleucine (purple). All carrying capacities are significantly different from each other unless otherwise marked (two way ANOVA with post-hoc Tukey's HSD, $p < 0.05$). N.S., not significant. Asterisk indicates a significant difference at the $p < 0.05$ level.

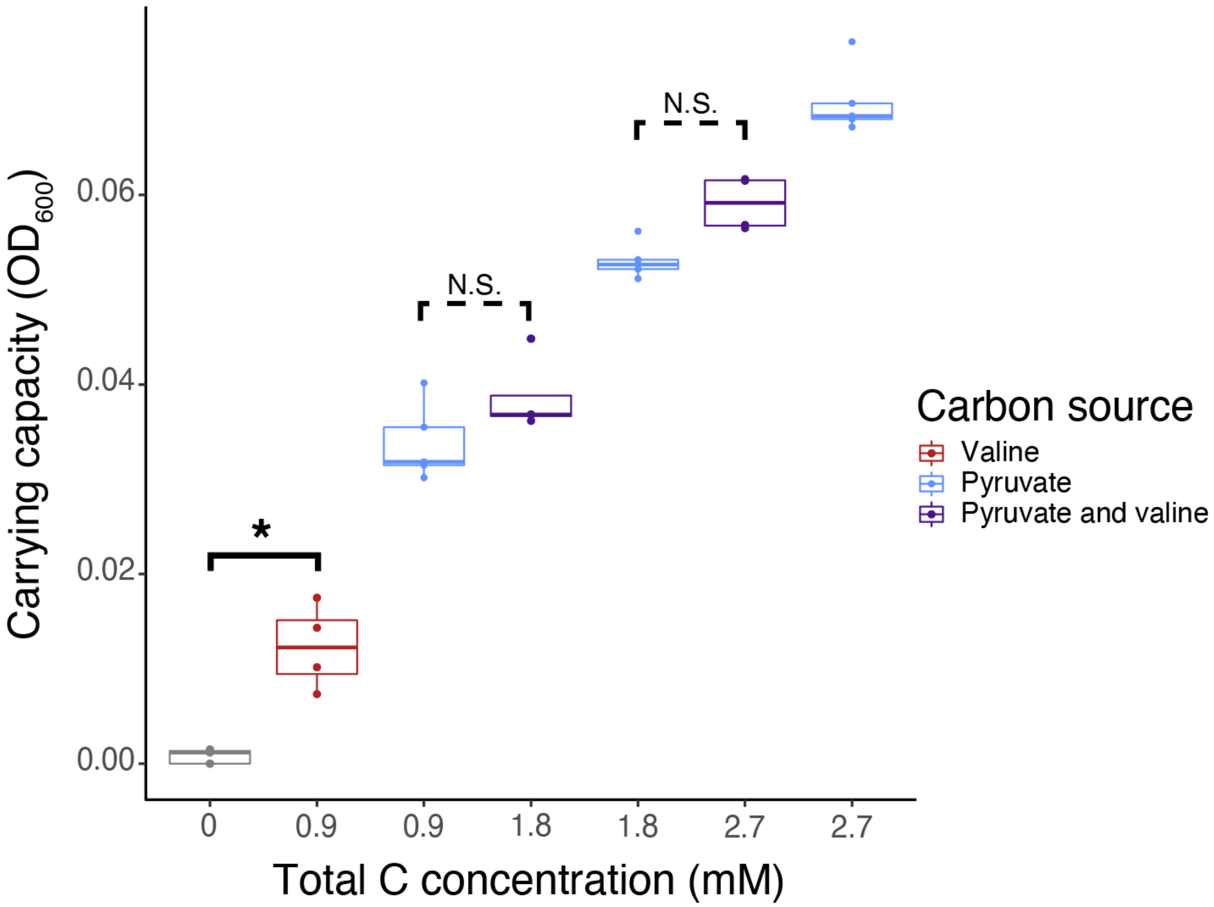


Figure S7. Calculated carrying capacities of *Alteromonas* incubated with valine (red), pyruvate (blue), or a mix of pyruvate and valine (purple). All carrying capacities are significantly different from each other unless otherwise marked (two way ANOVA with post-hoc Tukey's HSD, $p < 0.05$). N.S., not significant. Asterisk indicates a significant difference at the $p < 0.05$ level.

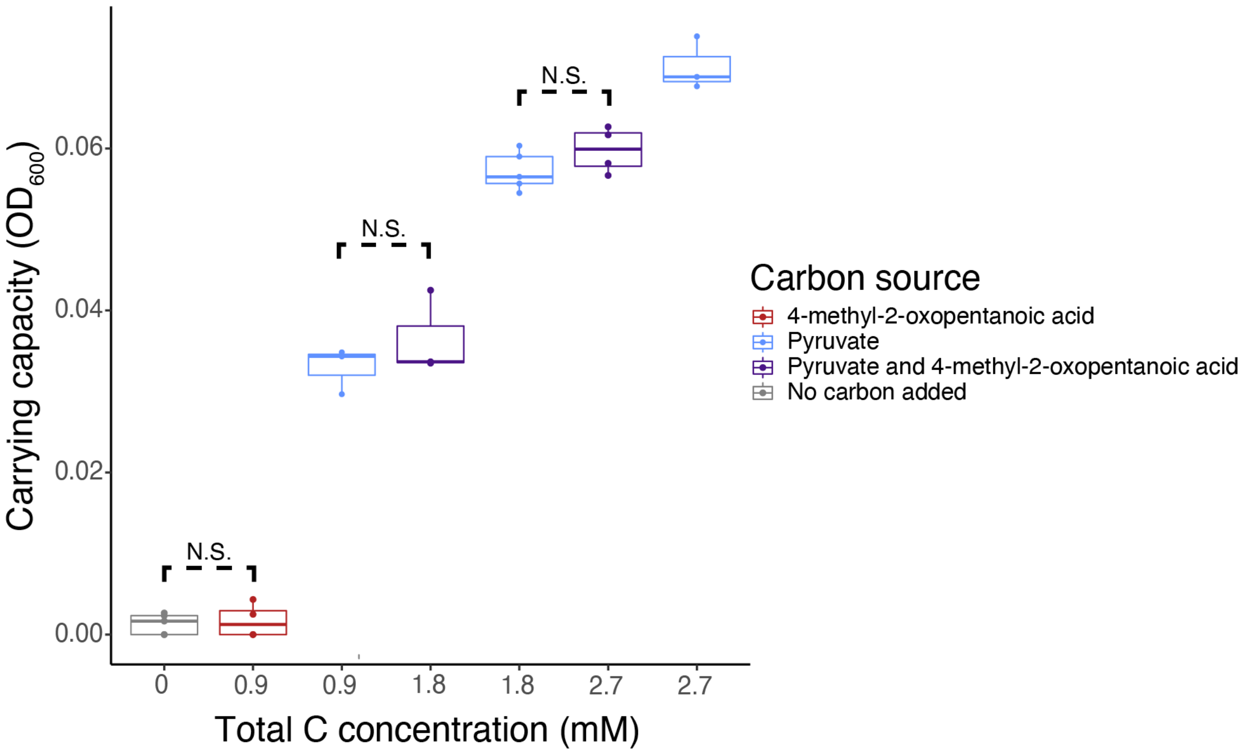


Figure S8. Calculated carrying capacities of *Alteromonas* incubated with 4-methyl-2-oxopentanoic acid (red), pyruvate (blue), or a mix of pyruvate and 4-methyl-2-oxopentanoic acid (purple). All carrying capacities are significantly different from each other unless otherwise marked (two way ANOVA with post-hoc Tukey's HSD, $p < 0.05$). N.S., not significant.

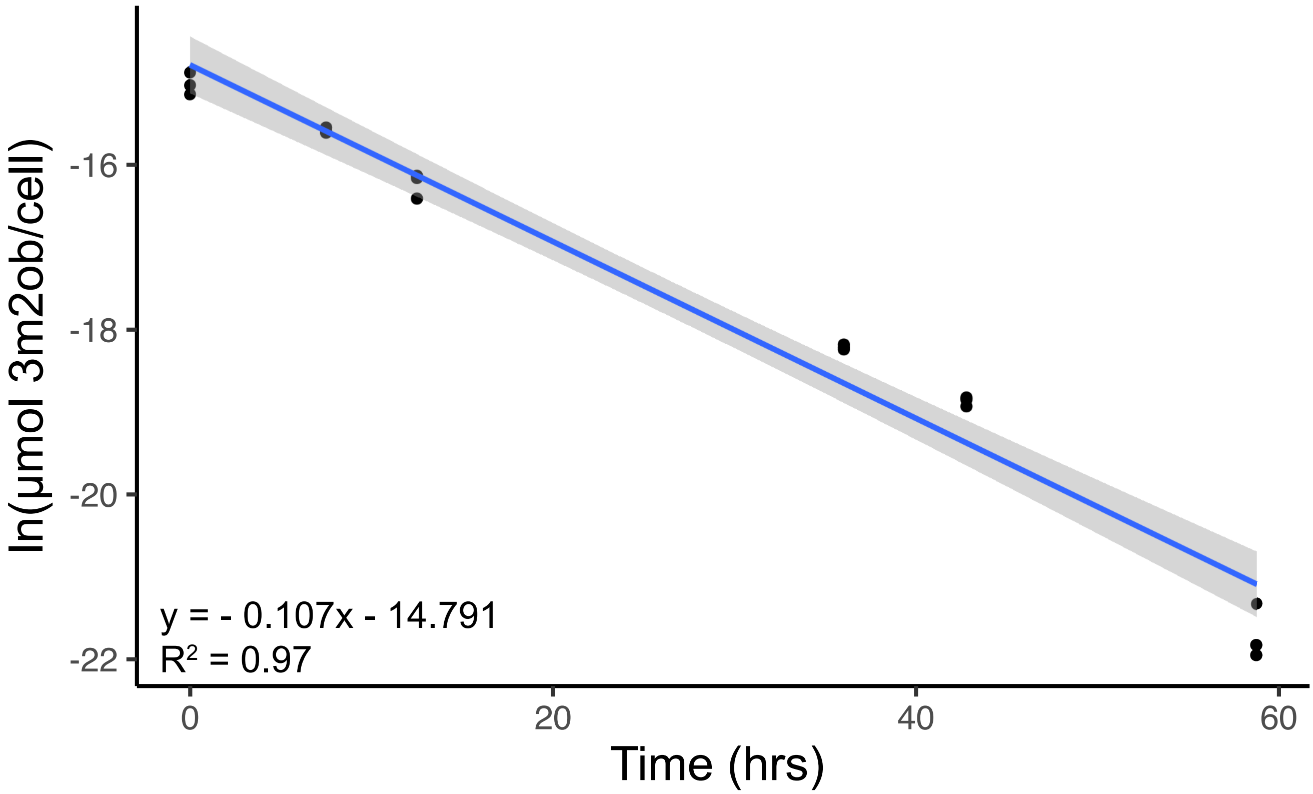


Figure S9. Cell-normalized concentrations of extracellular 3-methyl-2-oxobutanoic acid over the course of a 60 hour incubation with *Alteromonas macleodii* MIT1002.

#### Supplemental Works Cited


Agrawal, S., Kumar, S., Sehgal, R., George, S., Gupta, R., Poddar, S., Jha, A., & Pathak, S. (2019). El-MAVEN: A Fast, Robust, and User-Friendly Mass Spectrometry Data Processing Engine for Metabolomics. In A. D'Alessandro (Ed.), *High-Throughput Metabolomics* (Vol. 1978, pp. 301–321). Springer New York. https://doi.org/10.1007/978-1-4939-9236-2_19

Berube, P. M., Biller, S. J., Kent, A. G., Berta-Thompson, J. W., Roggensack, S. E., Roache-Johnson, K. H., Ackerman, M., Moore, L. R., Meisel, J. D., Sher, D., Thompson, L. R., Campbell, L., Martiny, A. C., & Chisholm, S. W. (2015). Physiology and evolution of nitrate acquisition in *Prochlorococcus*. *The ISME Journal*, *9*(5), 1195–1207. https://doi.org/10.1038/ismej.2014.211

Chambers, M. C., Maclean, B., Burke, R., Amodei, D., Ruderman, D. L., Neumann, S., Gatto, L., Fischer, B., Pratt, B., Egertson, J., Hoff, K., Kessner, D., Tasman, N., Shulman, N., Frewen, B., Baker, T. A., Brusniak, M.-Y., Paulse, C., Creasy, D., … Mallick, P. (2012). A cross-platform toolkit for mass spectrometry and proteomics. *Nature Biotechnology*, *30*(10), 918–920. https://doi.org/10.1038/nbt.2377

R Core Team. (2021). *R: A Language and Environment for Statistical Computing* [Computer software]. R Foundation for Statistical Computing. https://www.R-project.org/
